# Massively Multiplexed Affinity Characterization of Therapeutic Antibodies Against SARS-CoV-2 Variants

**DOI:** 10.1101/2021.04.27.440939

**Authors:** Emily Engelhart, Randolph Lopez, Ryan Emerson, Charles Lin, Colleen Shikany, Daniel Guion, Mary Kelley, David Younger

## Abstract

Antibody therapies represent a valuable tool to reduce COVID-19 deaths and hospitalizations. Multiple antibody candidates have been granted emergency use authorization by the FDA and many more are in clinical trials. Most antibody therapies for COVID-19 are engineered to bind to the receptor-binding domain (RBD) of the SARS-CoV-2 Spike protein and disrupt its interaction with ACE2. Notably, several SARS-CoV-2 strains have accrued mutations throughout the RBD that improve ACE2 binding affinity, enhance viral transmission, and escape some existing antibody therapies. Here, we measure the binding affinity of 33 therapeutic antibodies against a large panel of SARS-CoV-2 variants and related strains of clinical significance to determine epitopic residues, determine which mutations result in loss of binding, and predict how future RBD variants may impact antibody efficacy.

**One-Sentence Summary:** By measuring protein binding *in vitro*, we identify which clinical antibodies retain binding to various mutant SARS-CoV-2 strains.

## Introduction

Antibody therapies represent a valuable tool to reduce COVID-19 deaths and alleviate the burden on healthcare systems. During the vaccine rollout, antibody therapies to treat COVID-19 serve as a stopgap to save lives. Once vaccines are widely available, antibody therapies will continue to play an essential role in treating patients who are unvaccinated, such as the immunocompromised, and those infected with viral variants that escape vaccine protection. Antibody therapies can be developed quickly, making them well suited for rapid response to newly emerging strain variants, and have proven to reduce both viral loads and hospitalizations (*1, 2*). Multiple antibody candidates have been granted emergency use authorization by the FDA and many more are in phase 2 and phase 3 clinical trials (*3-5*).

Most antibody therapies for COVID-19 are engineered to bind to the receptor-binding domain (RBD) of the SARS-CoV-2 Spike protein and disrupt its interaction with angiotensin-converting enzyme 2 (ACE2) (*6-10*). Since the first human transmission of COVID-19 over one year ago, SARS-CoV-2 has undergone significant antigenic drift arising from mutations throughout the RBD that improve ACE2 binding affinity, enhance viral transmission, and generate resistance to existing antibody therapies (*11-16*). Understanding the impact of observed and likely RBD variants on the effectiveness of antibody therapies is of critical importance.

A handful of antibody therapies have been granted emergency use authorization by the FDA, over a dozen are in late-stage clinical development, and many more are in preclinical development (*17*). However, previous efforts to measure the effect of RBD variants on the efficacy of antibody candidates for COVID-19 have mostly been limited to characterizing individual CoV-2 RBD variants on a small number of antibody candidates (*8, 18-23*). A notable exception is the method introduced by Starr et al. which involves the construction of a yeast surface display library of RBD variants and enables the complete mapping of binding between a single antibody and all single mutations of the RBD (*14, 15, 24, 25*). The low antibody throughput of this method, however, precludes characterization of a wide variety of clinically relevant antibodies.

The AlphaSeq assay, previously described as yeast synthetic agglutination, is presented here as a method for overcoming existing throughput challenges for characterizing the binding profiles of tens of antibodies against thousands of RBD variants (*26*). Here, we leveraged the AlphaSeq assay to measure approximately 178,760 protein-protein interactions (PPIs) between 33 therapeutically relevant antibody candidates and most single-amino-acid mutations to SARS-CoV-2 RBD, along with selected widely circulating RBD variants containing multiple mutations. The PPI measurements are analyzed to derive key epitope residues for each antibody, determine individual or multiple mutations that result in loss of binding for each antibody, and predict how future RBD variants may impact antibody efficacy. **Figure 1** presents the AlphaSeq assay and its application to antibody/RBD binding in schematic form.

**Figure 1:**
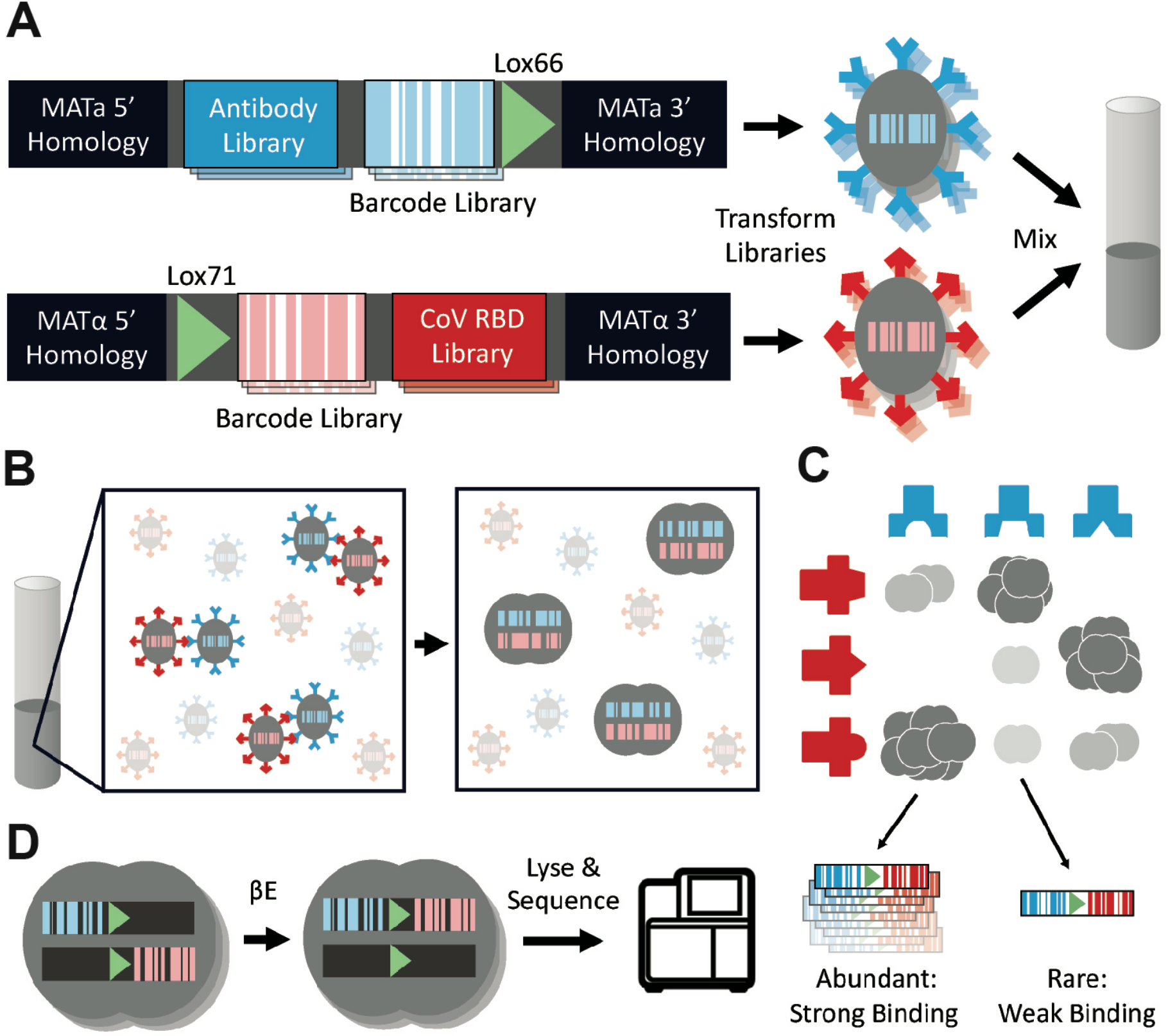
Using AlphaSeq to characterize interactions between an antibody library and a library of coronavirus RBD variants in high throughput. **A)** Two DNA fragment libraries are constructed with homology to the MATa or MATα genome for integration into the chromosome. Each library contains a diversity of proteins of interest, either antibodies or RBD variants, for display on the yeast cell surface, a diversity of randomized 25 nucleotide DNA barcodes, and a lox recombination site. MATa and MATα yeast strains lacking expression of native sexual agglutination proteins are transformed with their respective fragment library and subsequently mixed in liquid culture. **B)** In liquid culture, MATa-MATα agglutination is facilitated by interactions between surface displayed antibodies and RBD variants. Agglutination leads to mating between MATa and MATα haploid cells to produce a diploid cell. **C)** The number of diploids formed by a haploid pair is dependent on the interaction strength between the antibody and RBD variant expressed on their surfaces. **D)** Diploid cells are cultured with β-Estradiol to induce for CRE recombinase expression and recombine the engineered chromosome to pair DNA barcodes. Diploids are then lysed and sequenced to count the abundance of each barcode pair and determine the relative interaction strength between each antibody and coronavirus RBD variant.

## Results

First, to determine the epitope for each antibody an AlphaSeq experiment was performed where 33 antibodies were screened against a site-saturation mutagenesis (SSM) library comprising of (after library preparation and sequencing) approximately 75% of all single-residue mutants of 165 sites within the SARS-CoV-2 RBD **(Fig. 2A, S1, S2)**. While many RBD sites were intolerant to mutations (i.e., observed binding was poor across all antibodies and all mutants at that site), other sites revealed differential patterns of mutation-sensitivity among antibodies indicating epitopic diversity among the tested antibodies **(Fig. 2B)**. These sites allowed us to identify the epitope targeted by each antibody. Specifically, a pairwise binding comparison across all antibodies was carried out for each residue of the SARS-CoV-2 RBD **(Fig. 2C)**. For each site in the RBD included in the SSM library, all mutant affinities (scaled to each antibody’s affinity to WT RBD) were compared between the two antibodies with a Mann-Whitney U test (**Table S7**). For each antibody, sites at which differential sensitivity (Bonferroni-corrected p-values ≤ .05) was observed for at least 4 other antibodies were considered putative epitope residues (**Table S8**). Overall, 21 antibodies had residues that were found to have putative epitope residues **(Fig. 2D)**. For antibodies with known RBD-Ab structures, we found that our results correctly place key epitopic residues at the interface between the antibody and RBD **(Fig. 2E)**. Our results agree with previously published epitope data for COR-101, Imdevimab, Casirivimab, Bamlavimab, CC12.1 (*19, 27, 28*) while providing new insight into the epitopes targeted by the remaining antibodies and therefore allow us to prospectively identify which RBD sites should be of particular concern for antibody binding in new strains.

**Figure 2.**
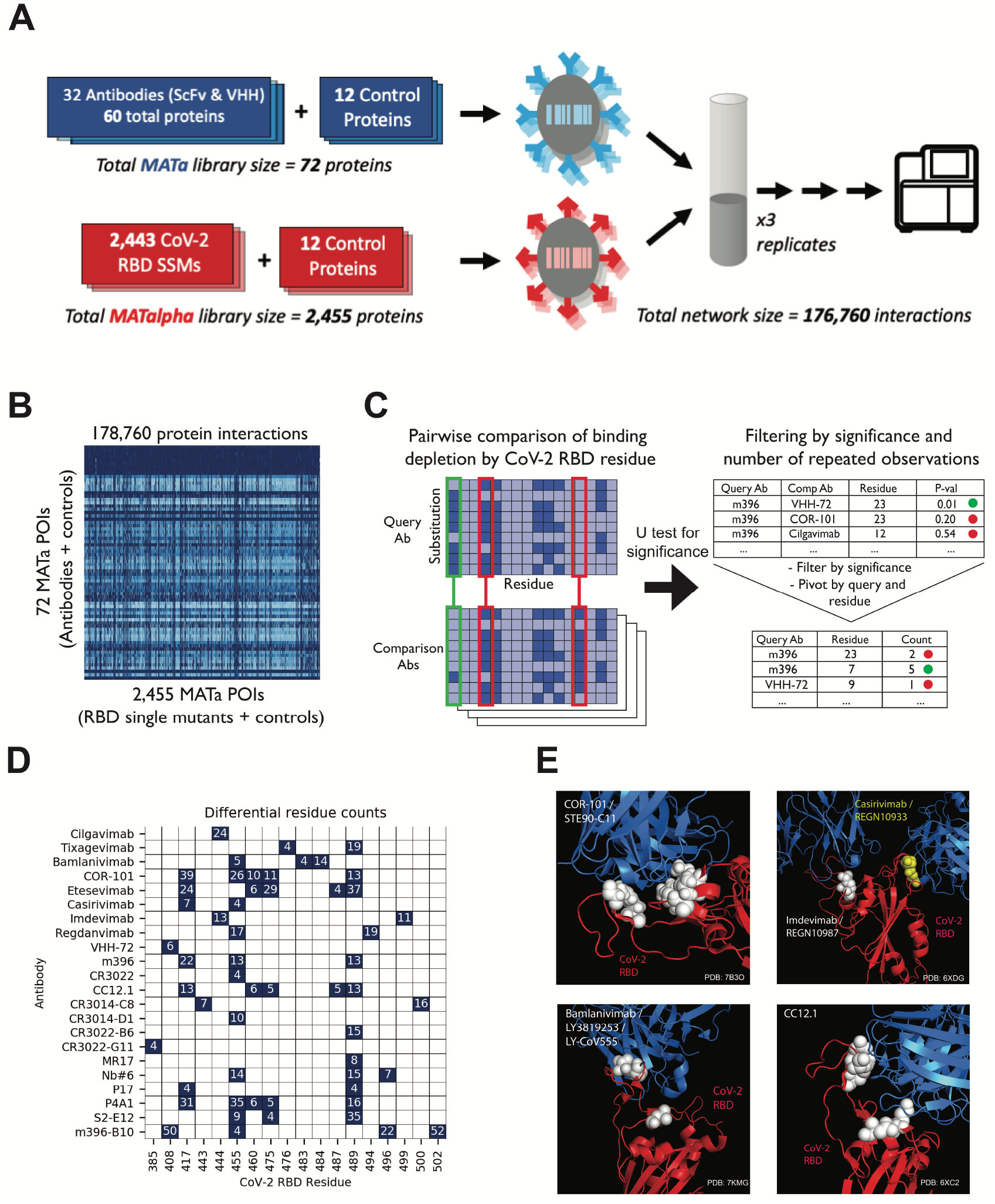
**A)** Schematic of epitope mapping AlphaSeq experiment, comparing antibodies against a CoV-2 RBD SSM library. **B)** Heatmap of AlphaSeq binding data showing all interactions measured in the assay in log10 KD (nM). RBD-CoV-2 intolerant substitutions results in loss binding for all antibodies in the set and appear as vertical dark blue streaks. **C)** Method summary for determination of epitope residues; each antibody was compared against all others, and at each RBD site a Mann-Whitney U test was performed to determine if binding was more impacted by a RBD mutations at that site in one antibody; results were filtered by U-test significance and for significance in 4 or more pairwise comparisons. **D)** Summary of epitope determination results –for each RBD site & antibody (summing results from LH and HL orientations where both were tested), # of pairwise comparisons with significant results. **E)** Representative antibodies (in blue) with known structure binding to RBD (in red), with residues called as epitope locations marked in white or yellow.

Next, a second AlphaSeq experiment was performed to assess binding of 33 clinically relevant antibodies and ACE2 against a curated panel of coronavirus RBD variants. Included were 34 unique SARS-CoV-2 RBD variants with single, double, or triple mutations, including 5 CDC-defined variants of concern, B.1.1.7, B.1.351, P.1, B.1.427, and B.1.429, and four related coronavirus RBDs **(Fig. 3A)**. From this single experiment, we determined the tolerance or enhancement of ACE2 binding, quantitatively measured the binding of each antibody to each SARS-CoV-2 variant and characterized the cross-reactivity profile of each antibody to related coronavirus strains **(Fig. S3, Table S9)**.

**Figure 3.**
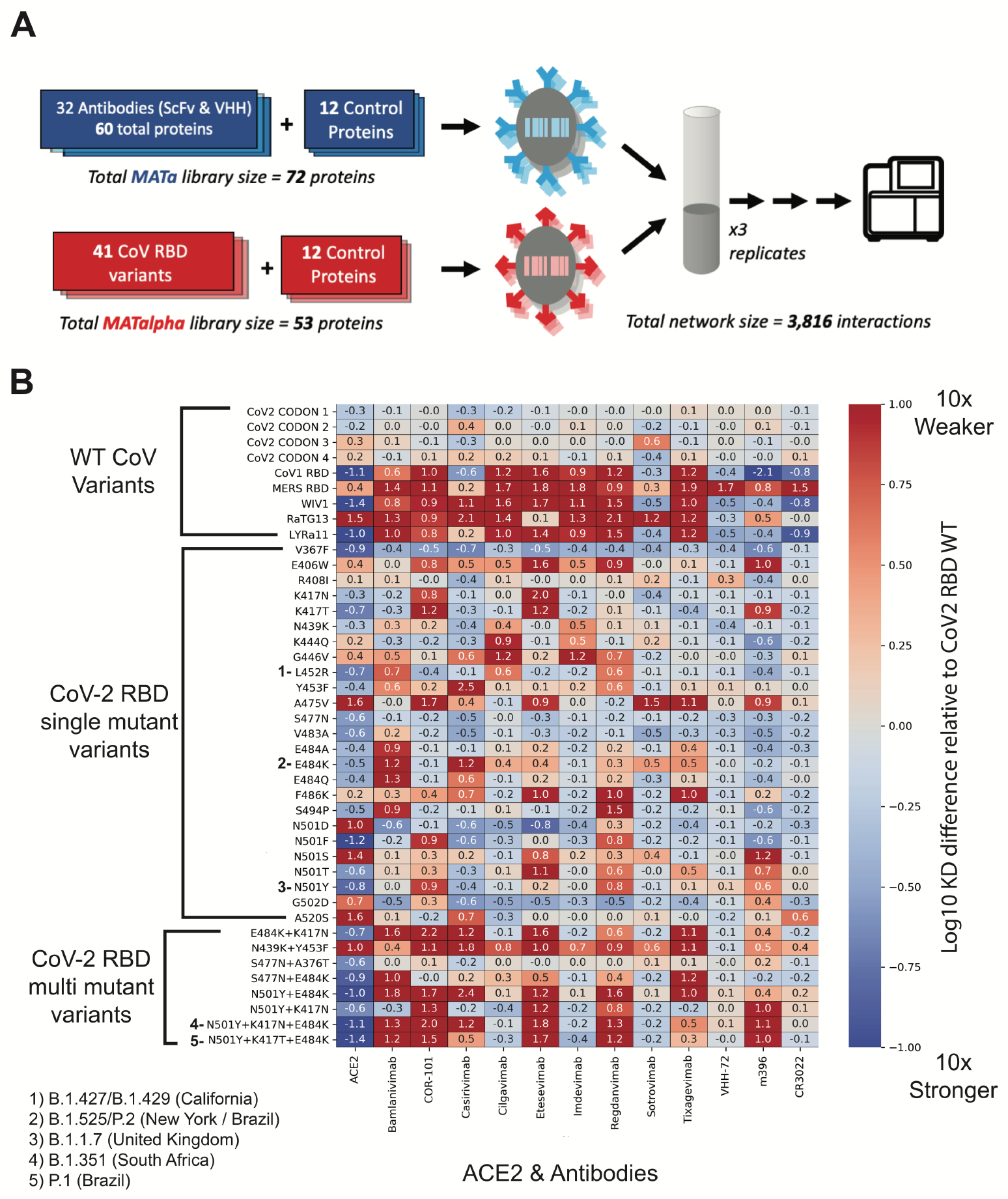
**A)** Schematic of AlphaSeq experiment, comparing antibodies against a curated panel of coronavirus RBD variants. **B)** Binding affinity of selected SARS-CoV-2 RBD variants to selected antibodies. Antibodies (or human ACE2) are on the x-axis, SARS-CoV-2 RBD variants are on the y-axis; in each column, values are mean difference in predicted binding affinity (measured on a log10 scale) between a given CoV variant and WT SARS-CoV-2 RBD for each antibody. Values below 0 (blue) represent improved binding relative to WT SARS-CoV-2 RBD and values above 0 (red) represent reduced binding affinity. In these data, Cilgavimab/AZD1061, Imdevimab/REGN10987 and Sotrovimab/GSK4182136 retain high affinity for the widest variety of clinically relevant RBD variants.

Measuring the interactions with ACE2 revealed many SARS-CoV-2 RBD variants with tighter receptor binding. In particular, all 5 of the CDC’s variants of concern showed increased binding affinity to ACE2, which is supported by previously reported affinities for B.1.1.7 and B.1.351 (*29*) and molecular dynamics simulations for B.1.1.7, B.1.351, and P.1 (*30*).

The dataset includes 9 antibodies in clinical trials with publicly available sequences (**Table S2)** (*31*). We observed a wide range of binding to the panel of SARS-CoV-2 RBD variants (**Fig. 3B**). Notably, we found good concordance with previous literature reports for most antibody-variant interactions **(Table S1)**. Of these 9 antibodies, 3 antibodies have been extensively studied in the literature: Imdevimab, Casirivimab and Bamlanivimab (*14, 18, 22, 23, 25, 32, 33*). We recapitulated the binding profiles for the B.1.1.7, B.1.351, and P.1 variants to Imdevimab, Casirivimab and Bamlanivimab (*23, 32, 33*). To summarize: we observe no significant changes in binding to the B.1.1.7 variant for Imdevimab, Casirivimab or Bamlanivimab. We observed a decrease in binding affinity for Casirivimab and Bamlanivimab to B.1.351 and P.1 variants whereas Imdevimab retained WT binding. Our findings with these 4 antibodies are highly consistent with previous literature reports. To date, there have been minimal or no reports on Sotrovimab, Regdanvimab, Tixagevimab, Cilgavimab, and COR-101 antibodies (*19, 24*). We find that Regdanvimab and COR-101 display reduced binding affinity to B.1.1.7, B.1.351, and P.1 variants. Tixagevimab has reduced binding affinity to B.1.351, and P.1 variants but maintains WT binding affinity with the B.1.1.7 variant. Whereas Sotrovimab and COR-101 show no changes in binding affinity to the B.1.1.7, B.1.351, and P.1 variants.

As part of the same experiment, cross-reactivity was evaluated by measuring antibody binding to four additional RBDs from related coronaviruses: SARS-CoV-1, LYRa11, WIV1, and RaTG13. Some antibodies were found to be highly specific to SARS-CoV-2, including Imdevimab, Bamlanivimab, Regdanvimab, Tixagevimab, Cilgavimab, and COR-101. In contrast, Casirivimab, Sotrovimab, and Etesevimab demonstrate varying degrees of cross-reactivity.

From two AlphaSeq assays, we mapped binding of ACE2 and 33 CoV antibodies against a panel of SARS-CoV-2 RBD variants, including B.1.1.7, B.1.351, and P.1, and identified key epitope residues for each antibody. This experiment has provided for the first time a comprehensive view of the impact of potential escape mutants (including several strains of substantial clinical concern) on the binding of RBD-targeted antibody therapeutics. For the small fraction of the dataset for which binding has been assayed previously, or for which the epitope is known via crystal structure, good agreement with previous methods establishes the reliability of AlphaSeq as a method. A full understanding of the impact of RBD mutations on the binding of therapeutic antibodies is essential to prioritize the development of therapies effective against the widest possible variety of circulating and newly emerging viral variants, and to enable the eventual possibility of regional or even personal prioritization of the most effective therapies for circulating SARS-CoV-2 variants.

## Supporting information

Supplementary Table 1 - Literature Comparison

Supplementary Table 2 - Antibody Sequences

Supplementary Table 3 - RBD & ACE2 Sequences

Supplementary Table 4 - CoV2 SSM Raw Data

Supplementary Table 5 - CoV Curated Raw Data

Supplementary Table 6 - Antibodies Naming Format

Supplementary Table 7 - Epitope Comparison Raw Results

Supplementary Table 8 - Epitope Comparison Filtered Results

Supplementary Table 9 - Curated Fold Difference Results

## Acknowledgments

We thank Dr. Jesse Bloom and Tyler Starr for their help with antibody and RBD variant selection.

## Author contributions

Conceptualization: DY, RL, EE, RE. Formal analysis: RL, EE, RE. Funding acquisition: DY, RL. Investigation: EE, CL, MK, DG, CS. Methodology: RL. Project administration: RL, EE, MK. Supervision: RL. Visualization: RL, EE, RE. Writing – original draft: DY, RL, EE, RE. Writing – review & editing: DY, RL, EE, RE, CL

## Data and materials availability

Code is available at https://github.com/A-AlphaBio/cov2_antibodies_variants. Protein sequences are included in supplementary tables 2 & 3. Raw affinity data is available in supplementary tables 4 and 5.

## Supplementary Materials

Materials and Methods

Figs. S1 to S3

Tables S1 to S9

