## Supplementary Text for "Massively Multiplexed Affinity Characterization of Therapeutic Antibodies Against SARS-CoV-2 Variants"

**This PDF file includes:**

Materials and Methods
Figs. S1 to S3
Captions for Tables S1 to S9

**Other Supplementary Materials for this manuscript include the following:**

Table S1. (separate file) Literature Validation. Comparison between AlphaSeq derived values for curated RBD variants and antibodies compared to literature results.
Table S2. (separate file) Antibody Sequences. Data table containing a full description of the antibodies used in this work including sequencing information and source.
Table S3. (separate file) CoV Sequences. Data table containing the sequences for the wild-type CoV RBD truncations used in this work.
Table S4. (separate file) Raw binding measurements for the epitope mapping assay.
Table S5. (separate file) Raw binding measurements for curated CoV variants assay.
Table S6. (separate file) Antibody naming convention.
Table S7. (separate file) Antibody pairwise CoV-2 substitution comparisons.
Table S8. (separate file) Epitope results summary.
Table S9. (separate file) CoV-2 WT normalized binding data.

### Materials and Methods

#### Materials

Antibody candidates

Clinically relevant antibodies were identified based on those designated to be in clinical trials in the COVID-19 Antibody Therapeutics tracker on January 9th, 2021 (*1*) Nine antibody candidates were found to have amino acid sequences publicly available at the time after a literature search and were included in this work: REGN10933 (casirivimab), REGN10987 (imdevimab), Bamlanivimab (LY3819253, LY-CoV555), Regdanvimab (CT-P59), Sotrovimab (VIR-7831/GSK4182136), AZD8895/Tixagevimab/COV2-2196, AZD1061/Cilgavimab, Etesevimab, COR-101 / STE90-C11. Additional antibodies were identified from the literature and their sequences were obtained from the Coronavirus-Binding Antibody Sequences & Structures (CoV-AbDab) (*2*) Overall, a total of 33 distinct antibodies were selected for this work (27 IGG's and 6 VHH formats). IGG format antibodies were built as single chain variable fragments (scFvs) in both heavy-light and light-heavy orientations. More detailed information for each antibody can be found in Supplementary Table #2.

Coronavirus variants

SARS-CoV-2 (isolate Wuhan-Hu-1, Genbank accession number MN908947, residues 319-527) and additional sarbecovirus homologs (RaTG13, Genbank MN996532; SARS-CoV-1 Urbani, Genbank AY278741; WIV1, Genbank KF367457; LYRa11, Genbank KF569996). CoV RBD sequences can be found in Supplementary Table #3.

Yeast media

Yeast peptone dextrose (YPD), yeast peptone galactose (YPG), and synthetic drop out (SDO) media supplemented with 80 mg/L adenine were made according to standard protocols. Suppliers used for our yeast media are as follows: Bacto Yeast Extract (Life Technologies), Bacto Tryptone (Fisher BioReagents), Dextrose (Fisher Chemical), Galactose (Millipore Sigma), Adenine (ACROS Organics), Yeast Nitrogen Base w/o Amino Acids (Thermo Scientific), SC-His-Leu-Lys-Trp-Ura Powder (Sunrise Science Products), L-Histidine (Fisher BioReagents), L-Tryptophan (Fisher BioReagents), Uracil (ACROS Organics), and Bacto Agar (Fisher BioReagents).

#### Methods

Isogenic yeast plasmid transformation

AlphaSeq compatible plasmids encoding yeast surface display cassettes were constructed by Twist Bioscience and resuspended at 100ng/µL. 100ng of plasmid was digested with PmeI enzyme for 1hr at 37°C to linearize, leaving chromosomal homology for integration into the ARS314 locus at both the 5' and 3' ends as described in (*3*). Yeast transformations were performed with Frozen-EZ Yeast Transformation Kit II (Zymo Research) according to manufacturer's instructions. Yeast were plated on SDO-Trp plates and grown at 30°C for 2-3 days. Successful transformants were struck out onto YPAD plates and grown overnight at 30°C.

Isogenic yeast fragment transformation

AlphaSeq compatible fragments encoding yeast surface display cassettes were constructed by Twist Bioscience and resuspended at 100ng/µL. A 3 piece yeast transformation was performed with upstream and downstream fragments which contained chromosomal homology for integration into the ARS314 locus at both the 5' and 3' ends as described in (*3*). Yeast transformations were performed with Frozen-EZ Yeast Transformation Kit II (Zymo Research) according to manufacturer's instructions. Yeast were plated on SDO-Trp plates and grown at 30°C for 2-3 days. Successful transformants were struck out onto YPAD plates and grown overnight at 30°C.

Protein expression validation – Flow cytometry

Yeast was inoculated in YPAD and grown overnight at 30°C. Yeast were labelled with FITC-anti-C-myc antibody (Immunology Consultants Laboratory, Inc.) in PBS (Gibco) + 0.2% BSA (Thermo) for 30 minutes at RT. Yeast were pelleted and resuspended in PBS + 0.2% BSA and read on a LSRII cytometer.

DNA library construction

AlphaSeq compatible fragments were synthesized by Twist Bioscience and were resuspended at 1 ng/µL in molecular grade water and pooled together. Fragment libraries were PCR amplified using KAPA DNA polymerase (Roche). A second DNA fragment with a randomized DNA barcode was PCR amplified. Fragments were run on a 0.8% agarose gel and extracted using Monarch Gel Purification kit (NEB).

SSM library construction

SSM library of SARS-CoV-2 RBD was synthesized by Twist Bioscience and resuspended at 1 ng/µL in molecular grade water. The SSM library fragments were PCR amplified using KAPA DNA polymerase (Roche). qPCR was terminated before saturation to minimize PCR bias, generally between 12-15 cycles. A second DNA fragment with a randomized DNA barcode was PCR amplified. Fragments were run on a 0.8% agarose gel and extracted using Monarch Gel Purification kit (NEB).

Yeast library transformation

MATa or MATalpha AlphaSeq yeast were grown for 6 hours in YPAG media to induce SceI expression, as described in [Younger, et al.]. All spin steps were performed at 3000 RPM for 5 minutes. Yeast was spun down and washed once in 50 mL 1M Sorbitol (Teknova) + 1mM $CaCl_2$ solution. Washed yeast were resuspended in a solution of 0.1M LiOAc/1mM DTT and incubated shaking at 30°C for 30 minutes. After 30 minutes, yeast was spun down and washed once in 50 mL 1M Sorbitol + 1mM $CaCl_2$ solution. Yeast was resuspended to a final volume of 400 µL in 1M Sorbitol + 1mM $CaCl_2$ solution and incubated with DNA for at least 5 minutes on ice. Yeast were electroporated at 2.5kV and 25 uF (BioRad). Immediately following electroporation, yeast were resuspended in 5 mL of 1:1 solution of 1M Sorbitol:YPAD and incubated shaking at 30°C for 30 minutes. Recovered yeast cells were spun down and resuspend in 50 mL of SDO-Trp media and transferred to a 250mL baffled flask. 20 µL of resuspended cells were plated on SDO-Trp to determine transformation efficiency. Both the flask and plate were incubated at 30°C for 2-3 days. After 2-3 days, transformation efficiency was determined by counting colonies on the SDO-Trp plate.

Nanopore barcode mapping

Genomic DNA from yeast libraries was extracted using Yeast DNA Extraction Kit (Thermo Scientific) following the manufacturer's instructions. A single round of qPCR was performed to amplify a fragment pool from the genomic DNA containing the gene through the associated DNA barcode. qPCR was terminated before saturation to minimize PCR bias, generally between 15-20 cycles. The final amplified fragment was concentrated with KAPA beads, quantified with a Quantus (Promega), prepped with a SQK-LSK-110 ligation kit (Oxford Nanopore) and sequenced with a Minion R10 flow cell (Oxford Nanopore) following the manufacturer's instructions.

Library-on-Library AlphaSeq Assays

2 mL of saturated MATa and MATalpha library were combined in 800 mL of YPAD media and incubated at 30°C in a shaking incubator. 3 technical replicates were performed for each assay. After 16hr, 100 mL of yeast culture was washed once in 50 mL of sterile water and transferred to 600 mL of SDO-lys-leu with 100 nM ß-estradiol (Sigma) for 24hr. After 24hr, 100 mL of yeast was transferred to fresh SDO-lys-leu with 100 nM ß-estradiol for an additional 24hr.

Library preparation for Next-Generation Sequencing

Genomic DNA was extracted using Yeast DNA Extraction Kit (Thermo Scientific) following manufacturer's instructions. qPCR was performed to amplify a fragment pool from the genomic DNA and to add standard Illumina sequencing adaptors and assay specific index barcodes. qPCR was terminated before saturation to minimize PCR bias, generally between 23-27 cycles. The final amplified fragment was concentrated with KAPA beads, quantified with a Quantus (Promega), and sequenced with an NextSeq 500 sequencer (Illumina).

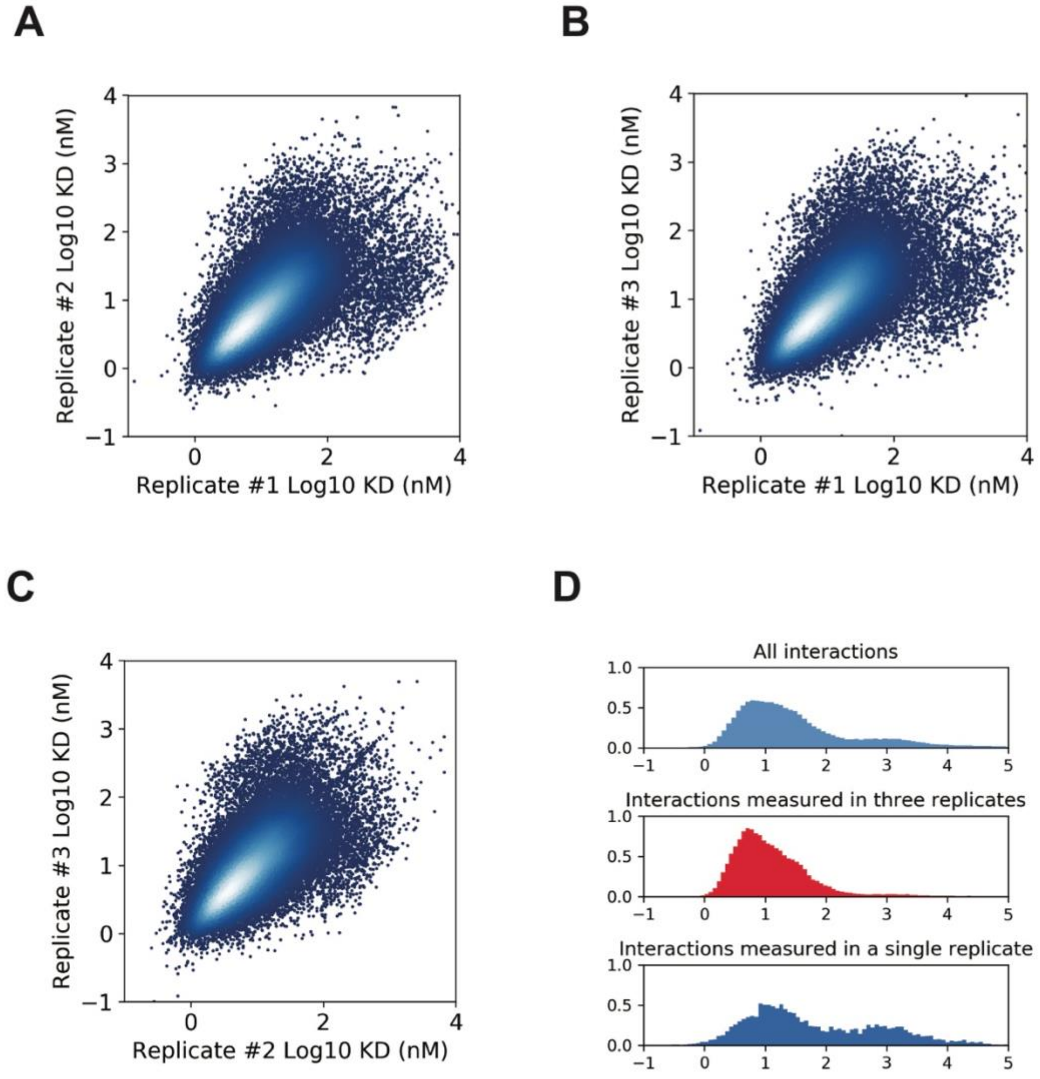


**Fig. S1. Overview of experimental reproducibility and binding affinity distribution for epitope mapping AlphaSeq experiment consisting of 178,760 interactions. A)** Scatter plot corresponding to the binding affinity measurements in replicate #1 and replicate #2 for interactions observed in both replicates. **B)** Scatter plot corresponding to the binding affinity measurements in replicate #1 and replicate #2 for interactions observed in both replicates. **C)** Scatter plot corresponding to the binding affinity measurements in replicate #2 and replicate #3 for interactions observed in both replicates. **D)** Binding affinity measurement distributions for all interactions, interactions observed in three replicates and interactions only observed in one replicate.

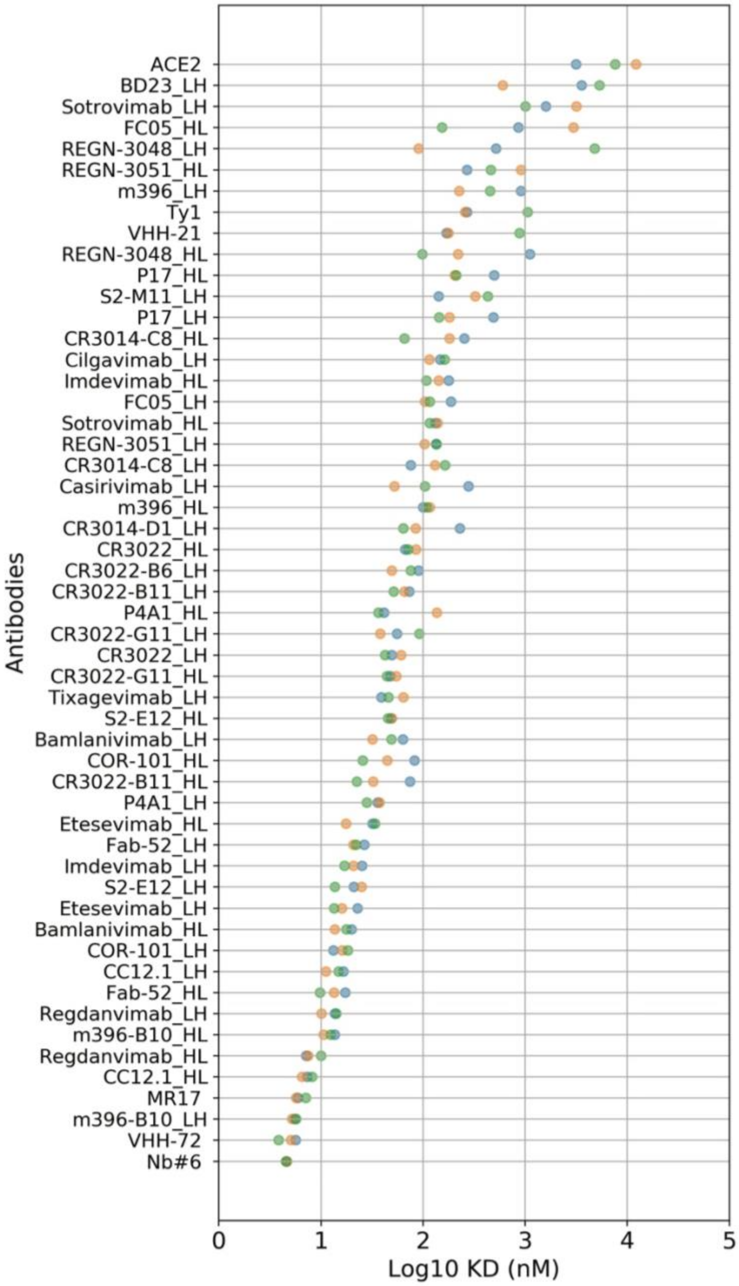


**Fig. S2. Interactions between wild-type SARS-CoV-2 RBD and binders observed in all three replicates in epitope mapping AlphaSeq assay.**

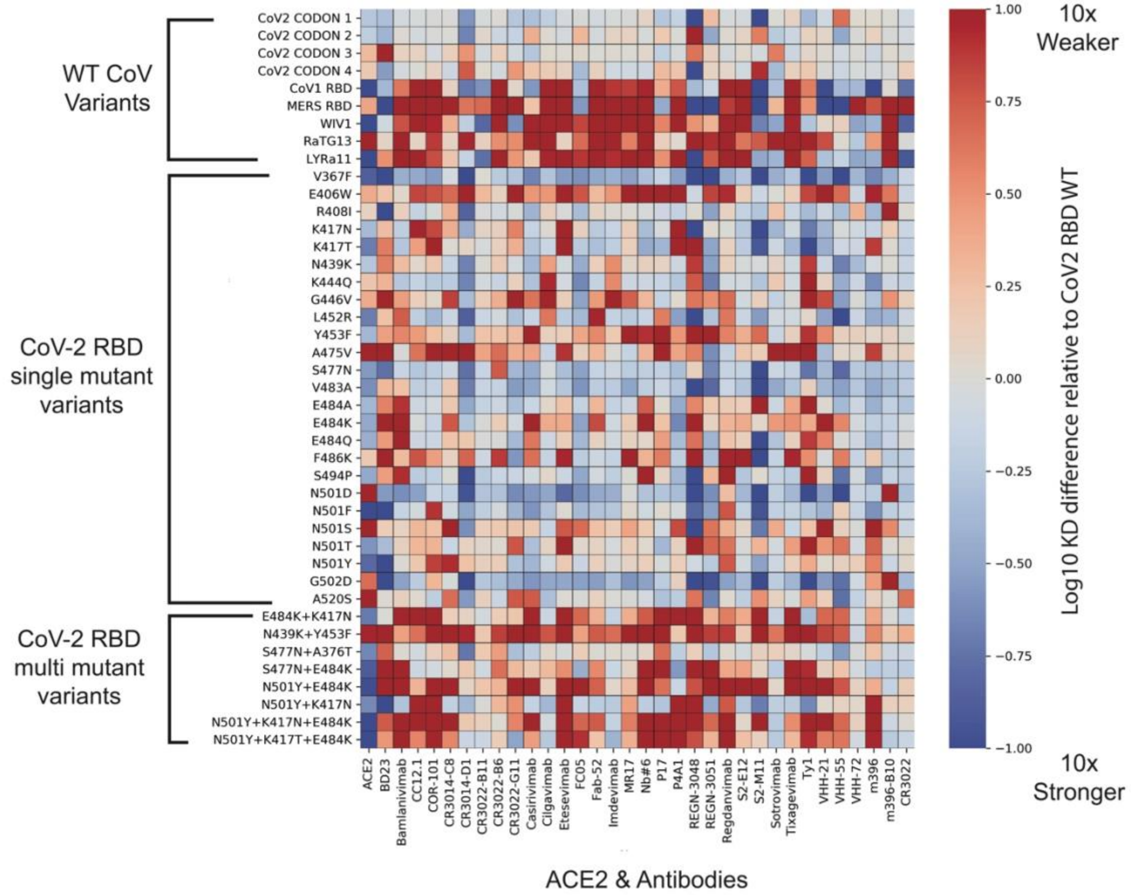


**Fig. S3. Binding affinity of selected SARS-CoV-2 RBD variants to all antibodies.**
Antibodies (or human ACE2) are on the x-axis, SARS-CoV-2 RBD variants are on the y-axis; in each column, values are mean difference in predicted binding affinity (measured on a log10 scale) between a given CoV variant and WT SARS-CoV-2 RBD for each antibody. Values below 0 (blue) represent improved binding and values above 0 (red) represent reduced binding affinity.

**Table S1. (separate file) Literature Validation.** Comparison between AlphaSeq derived values for curated RBD variants and antibodies compared to literature results. Overall, we found a very strong agreement between our reported affinity values for the subset of interactions that have been reported by other groups.

**Table S2. (separate file) Antibody Sequences.** Data table containing a full description of the antibodies used in this work including sequencing information and source.

**Table S3. (separate file) CoV Sequences.** Data table containing the sequences for the wild-type CoV RBD truncations used in this work.

**Table S4. (separate file) Raw binding measurements for the epitope mapping assay.** Matalpha_description field contains SSM descriptors with the following notation: wild type amino acid, amino acid position, amino acid substitution and codon sequence (e.g., V165Q_CAA). Amino acid position in the SSM library has an offset of 318 amino acids relative to the full length CoV-2 RBD. Pred_aff field contains log 10 KD affinity measurements as generated from the AlphaSeq assay after extrapolation from a standard curve with known affinities spiked into the assay. If a mating was not observed, the field is left blank, and it is presumed to be below the limit of detection for the assay. Sample_name corresponds to experimental mating replicates 1, 2 and 3.

**Table S5. (separate file) Raw binding measurements for curated CoV variants assay.** Pred_aff field contains log 10 KD affinity measurements as generated from the AlphaSeq assay after extrapolation from a standard curve with known affinities spiked into the assay. If a mating was not observed, the field is left blank, and it is presumed to be below the limit of detection for the assay. Sample_name corresponds to experimental mating replicates 1, 2 and 3.

**Table S6. (separate file) Antibody naming convention.**

**Table S7. (separate file) Antibody pairwise CoV-2 substitution comparisons.** Dataset containing all pairwise comparisons among antibodies in the set for binding sensitivity against SARS-CoV-2 RBD substitutions. Field 'diff' refers to the log10 KD affinity difference in binding between the 'query' antibody and the 'comparison antibody'.

**Table S8. (separate file) Epitope results summary**. Summary table of epitope determination results –for each RBD site & antibody (summing results from LH and HL orientations where both were tested), # of pairwise comparisons with significant results.

**Table S9. (separate file) CoV-2 WT normalized binding data.** Complete dataset of binding affinity of CoV variants to selected antibodies after normalization against binding for wild-type CoV-2 RBD. Values are mean difference in predicted binding affinity (measured on a log10 scale) between a given CoV variant and WT SARS-CoV-2 RBD for each antibody. Values below 0 (blue) represent improved binding and values above 0 (red) represent reduced binding affinity
